# Gradients in excitability generate hippocampal waves and shape their interactions with cortex

**DOI:** 10.64898/2026.05.27.728116

**Authors:** Anna Behler, Richa Phogat, David T. Jones, James M. Shine, Michael Breakspear

**Affiliations:** School of Science, College of Engineering, Science and the Environment, University of Newcastle, Callaghan, NSW, Australia; Brain Health Research Program, Hunter Medical Research Institute, Newcastle, NSW, Australia; Department of Neurology, Mayo Clinic, Rochester, MN, USA; School of Medical Sciences, The University of Sydney, Sydney, NSW, Australia

**Keywords:** hippocampus, travelling waves, cortical embedding, hippocampal-cortical interactions, neural mass models

## Abstract

Travelling waves are a prominent feature of hippocampal activity, but the mechanisms determining their propagation and influence on the cerebral cortex remain unclear. Using a biophysically-grounded model of neural activity evolving across hippocampal and cortical surfaces, we show that spatial gradients in external input or neural excitability are a sufficient mechanism for the emergence of travelling waves along the long axis of the hippocampus. These waves emerge only above a critical gradient threshold, propagate with biologically-plausible velocities and exhibit frequency-dependent reversals in direction across slow and fast theta regimes. When coupled to the cerebral cortex, anterior-to-posterior hippocampal waves selectively reorganise globally synchronous cortical activity into metastable travelling waves, whereas posterior-to-anterior hippocampal waves do not, revealing a directional asymmetry in hippocampal–cortical communication. Conversely, cortical waves aligned with large-scale functional gradients induce structured hippocampal waves, revealing complementary direction specific effects across the two systems. These analyses identify structured excitability gradients as a principle governing wave propagation in the hippocampus and suggest how wave-to-wave interactions may coordinate complex cortical and hippocampal interactions during mnemonic and perceptual processes.

## 1 Introduction

The hippocampus is a core neural hub that supports cognitive function through reciprocal exchange with large-scale cortical networks. This process requires integration of hippocampal activity into the complex dynamics of diverse cortical regions (Barnett et al., 2021; Ferris et al., 2025; Genon et al., 2021; Olsen et al., 2012). Behavioural profiling suggests that the hippocampus possesses a dominant organizational principle — or primary gradient — along its anterior-posterior (A-P) axis (Poppenk et al., 2013), such that anterior hippocampus supports self-centric processes, such as semantic, autobiographical and contextual associations, while posterior hippocampus preferentially supports world-centric processes including spatial and goal-directed perceptual processing, (Plachti et al., 2019; Vogel et al., 2020). This pattern of behavioural profiling is matched by gradients in microstructure and functional connectivity that are also aligned along the hippocampal long axis (Paquola et al., 2019; Przézdzik et al., 2019).

To achieve coordination with the cerebral cortex, these hippocampal gradients are topologically aligned with large-scale gradients of cortical function (Borne et al., 2023; Paquola et al., 2020). Accordingly, posterior hippocampus interacts with sensory cortex and goal-directed regions in the posterior cortex such as the parietal memory network (Zheng et al., 2021b), while more anterior hippocampal areas interact progressively with associative, then heteromodal and anterior default-mode networks (Barnett et al., 2021; Vos De Wael et al., 2018; Zheng et al., 2021a). Hippocampal and cortical dynamics thus mutually influence each other along this shared gradient, adapting to context through dynamic modulation of the underlying topographically organised projections (Borne et al., 2023). This shared principle supports the formation of coherent representations that integrate perceptual, conceptual, and mnemonic content (Olsen et al., 2012; Whittington et al., 2022).

Although the topological underpinning of this functional embedding has been established, the dynamics that coordinate hippocampal and cortical rhythms are not well understood. The organisation of neural activity into coherent travelling waves propagating across cortex is emerging as a fundamental principle of the brain’s complex dynamics (Ermentrout and Kleinfeld, 2001b; Muller and Destexhe, 2012; Roberts et al., 2019; Robinson et al., 1997). Travelling waves appear at different spatial scales and various states of consciousness (Massimini et al., 2004; Muller et al., 2016; Raut et al., 2021). Their spatiotemporal organisation indicates which cortical regions are currently active, and the direction in which that activity is propagating (Muller et al., 2018). Cortical output arises via interaction of these passing activity waves with local dendritic arborisations, particularly in superficial cortical layers (Heitmann et al., 2013). Accordingly, the direction and wavelength of propagating waves can encode specific cognitive (Xu et al., 2023)

and motor functions (Heitmann et al., 2012), including memory encoding and recall (Mohan et al., 2024; Zhang et al., 2018). Recent perturbational evidence further suggests that externally imposed travelling-wave dynamics can causally shape cognitive performance (Lee et al., 2026). Analyses of intracranial recordings suggests that low-frequency activity in the human hippocampus also forms travelling waves, propagating from the anterior head to the posterior tail (Lubenov and Siapas, 2009; Patel et al., 2012) and, in some cases, in the opposite direction (Kleen et al., 2021; Zhang and Jacobs, 2015). Hippocampal theta oscillations are a hallmark of these low-frequency rhythmic processes and play a central role in coordinating memory encoding, spatial navigation, and long-range communication with distributed cortical regions (Buzśaki, 2002; Colgin, 2016; Herweg et al., 2020). Rather than reflecting a unitary rhythm, theta comprises multiple functionally distinct components that may engage different cortical circuits depending on cognitive demands (Goyal et al., 2020). Therefore, the mechanisms generating hippocampal travelling waves and the conditions necessary for their emergence are of particular interest (Lubenov and Siapas, 2009; Patel et al., 2012).

Travelling theta waves may be a mechanism by which the hippocampus organises and integrates information from a wide array of neocortical networks. Bidirectional wave propagation along the A-P axis have been associated with memory consolidation processes on the cortical surface (Mohan et al., 2024) and in the hippocampus (Kleen et al., 2021). During sleep, slow hippocampal oscillations help coordinate both the expression and synchronisation of faster neocortical activities, suggesting a role for hippocampal waves in driving phase shifts that create travelling waves across the cortex (Cox et al., 2020).

While these empirical recordings offer intriguing insights into waves of hippocampal activity, their origin and functions remain unknown. Biophysically-informed computational modelling can address their origin and advance candidate principles for their functions. Within a computational framework, neural structures such as the hippocampus can be formalised as a coupled neural mass system (Deco et al., 2008), such that local dynamics are represented by locally interacting excitatory-inhibitory populations which then interact via long range projections. This framework allows the simulation of emergent phenomena, including theta-gamma nested oscillations, metastability and travelling waves (Hancock et al., 2025; Naoumenko and Gong, 2019; Roberts et al., 2019). Critically, hippocampal and cortical gradients can be derived from imaging data, ensuring that the model architecture is empirically constrained.

We study the emergent behaviour of coupled neural masses interacting on anatomically informed hippocampal and cortical surfaces. We first study the conditions supporting the emergence of waves propagating within the hippocampus. We benchmark the speed and wavelengths of these waves against those previously characterised in empirical data. We next study interacting cortical and hippocampal systems, with inter-system coupling informed by neuroimaging data acquired while human participants performed a naturalistic memory task. We thus investigate the reconfiguration of hippocampal and cortical activity under the influence of hippocampal to cortical and cortical to hippocampal coupling.

## 2 Results

### 2.1 Simulating cortical and hippocampal activity

Hippocampal-cortical activity was simulated using neural masses embedded within cortical and hippocampal surfaces (Fig. 1). Local neural populations comprise three interacting subpopulations (Jansen and Rit, 1995). Pyramidal cells project to local excitatory and inhibitory interneurons which each possess recurrent projections back to the pyramidal cells (Fig. 1A). In addition to these feedback loops, long-range pyramidal-to-pyramidal connections support interactions between neural masses across the hippocampal and cortical surfaces. Connection strengths within each surface decrease according to an exponential-distance rule (EDR), as widely observed in tracer (Horváat et al., 2016) and Magnetic Resonance Imaging (MRI)-based (Roberts et al., 2017) studies (Fig. 1B). Neural masses connected in this manner yield neural dynamics on both the cortex and hippocampus. We first study the ensuing isolated hippocampal dynamics (Section R2). Connections between hippocampus and cortex are then introduced (Fig. 1C, Suppl. Fig. S1), as constrained by functional neuroimaging data, enabling the study of cortical-hippocampal interactions (Section R3).

**Figure 1:**
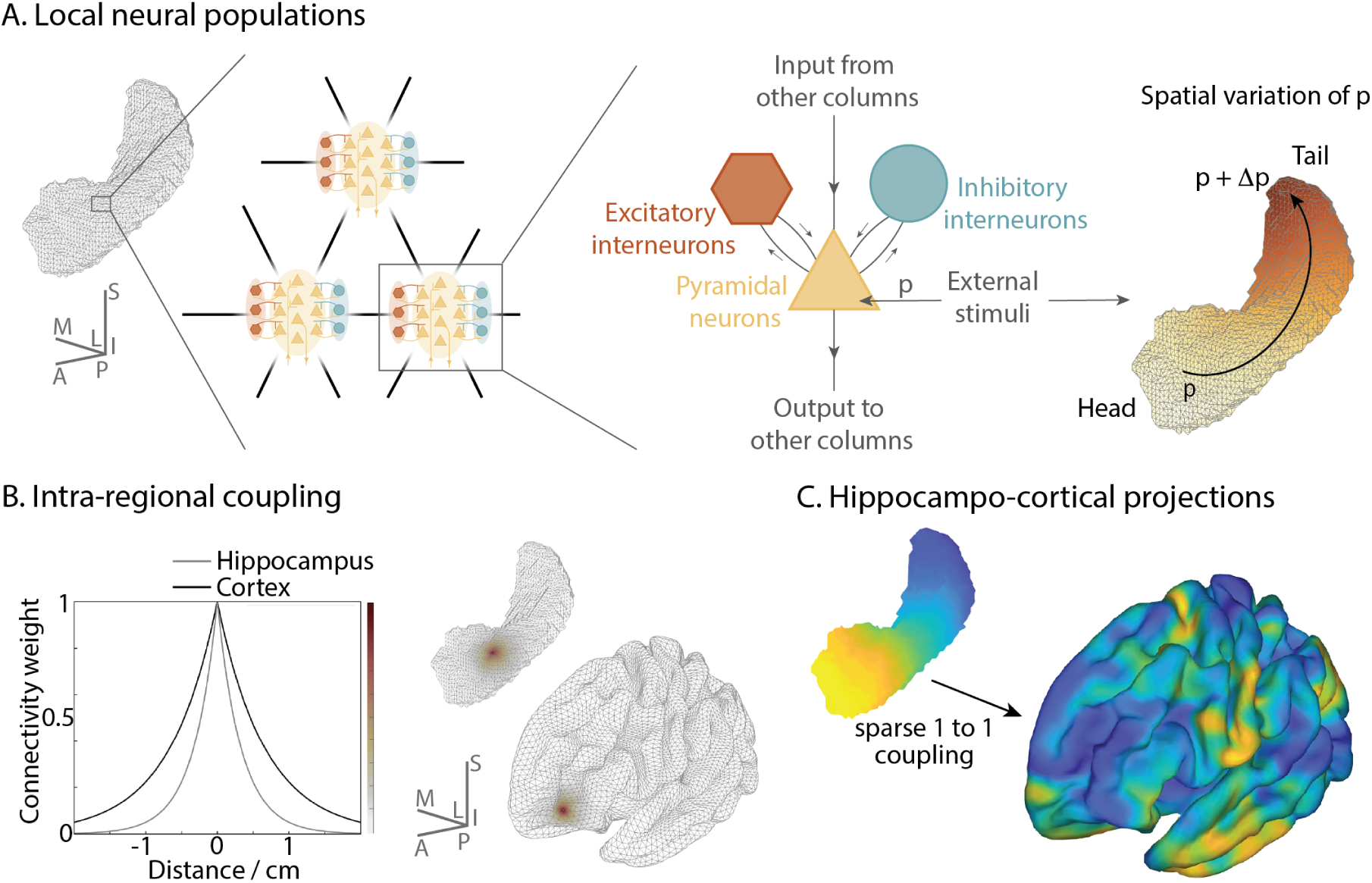
Implementation of neural mass models. **A** Jansen-Rit columns were placed at every vertex of hippocampal and cortical surface meshes. Each neural column consists of a population of pyramidal neurons with excitatory and inhibitory feedback loops. In addition to intra-column connections, pyramidal neurons project to distant columns and receive external inputs *p* as well as inputs from other Jansen-Rit columns. The external input *p* was spatially modulated along the primary AP-axis of the hippocampus. **B** Coupling between Jansen-Rit columns within the hippocampus and the cortex follow an exponential distance rule with exemplary values of the decay parameter *λ* (0.15 mm^—1^ for cortex and 0.30 mm^—1^ for hippocampus). **C** Hippocampus-to-cortex coupling was implemented as a sparse one-to-one mapping, where each hippocampal vertex is connected to exactly one cortical vertex (indicated by matching colours), informed by human functional connectivity data. See Methods for model equations and parameters.

### 2.2 External inputs and neuromodulation trigger hippocampal waves

We first focus on dynamics within the hippocampus, with the timescale and coupling constants of local populations chosen so that they exhibit either slow (4-7 Hz) or fast (5-9 Hz) theta oscillations (Table 1).

**Table 1:** Parameters for Jansen–Rit columns in the fast and slow theta regimes.

| Parameter | Fast theta | Slow theta |
| --- | --- | --- |
| $A$ | 5.6 | 7.0 |
| $B$ | 21 | 15 |
| $a$ ($\text{s}^{-1}$) | 80 | 100 |
| $b$ ($\text{s}^{-1}$) | 70 | 50 |
| $\bar{a}$ ($\text{s}^{-1}$) | 90 | 50 |
| $\alpha$ ($\text{s}^{-1}$) | 0.55 | 0.40 |
| $\beta$ ($\text{s}^{-1}$) | 0.35 | 0.25 |
| $p_{\text{head}}$ | 2.0 | 2.4 |

When the parameters of local populations are uniform across the hippocampus, the dynamics converge toward a globally synchronous state characterised by in-phase coherent theta oscillations (Fig. 2 and Suppl. Video 1). This is evident when viewing the population states colour-coded by their instantaneous phase (Fig. 2B, top) or by plotting the activity of local populations selected along the long axis of the hippocampus (Fig. 2B, bottom). The phase dynamics of all local populations approach complete alignment and the Kuramoto order parameter *R* which summarises the collective phase (see Methods) rapidly reaching its maximum value of 1 (Fig. 2A). This convergence to global synchrony with uniform parameter values occurs for both slow and fast theta oscillations.

**Figure 2:**
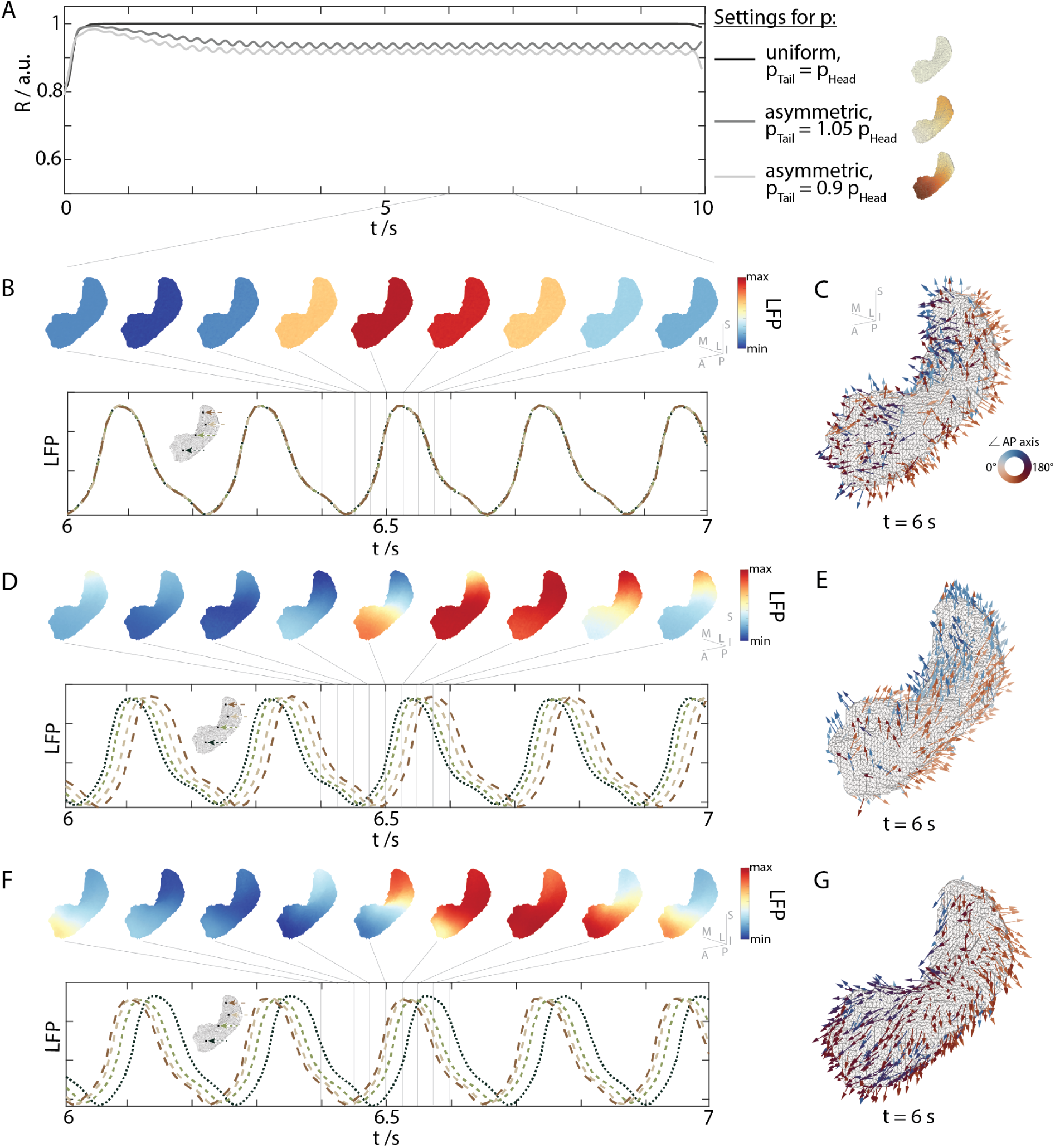
Self-organisation of hippocampal activity under spatially varying external input. *p* **across the anterior-posterior (A-P) axis. A** Global coherence *R* over time for each condition of spatially varying *p*. Uniform input yields near-total synchrony. Asymmetric inputs lead to reduced coherence, due to travelling wave dynamics. The accompanying colour bars show the spatial profile of *p* for each regime. **B** Local field potential (LFP) snapshots show homogenous states forming, as under uniform input (*p*_tail_ = *p*_head_) local populations rapidly synchronise, yielding globally coherent oscillations. LFP timeseries from five representative vertices across the A-P axis exhibit near-identical, overlapping oscillations. **C** Orientations of nodal phase velocity vectors are disorganised, indicating the absence of any wave activity. **D** Spontaneous emergence of travelling waves, after introducing a modest spatial gradient (*p*_tail_ *> p*_head_), visible in LFP snapshots. LFPs show consistent temporal lags across the sampled vertices. **E** Nodal velocity vectors align with the long hippocampal axis reflecting propagation of a wavefront from head to tail, in the opposite direction of the input current. **F** LFP patterns indicate reversed direction of wave propagation following reversing and slightly increasing the asymmetry (*p*_head_ *> p*_tail_). LFP time series maintain phase offsets, but in reverse order. **G** Phase vectors align tail to head confirming the directionality of the wave. See Suppl. Videos 1–5

We then introduced a spatially varying external input *p* to the hippocampus, maximum at the tail of the hippocampus and decreasing linearly along the P-A axis (Fig. 1 A). In the slow theta regime, when the ensuing imbalance is greater than 2% of the total current, hippocampal activity spontaneously reorganises into travelling waves that traverse from the head to the tail of the hippocampus, hence in the reverse direction of the input current gradient (Suppl. Video 2). This is evident in the emergence of a smooth gradation of phases along the hippocampal surface (Fig. 2 D, top) and likewise a successive phase delay of time series from local populations situated incrementally along the hippocampal axis (Fig. 2D, bottom). The duration of the initial transient before the reorganisation into waves depends on the magnitude of the imbalance: larger differences in the current between the tail and head led to a faster emergence of travelling waves. The temporal evolution of the Kuramoto order parameter captures the partial phase alignment within the wave patterns, reflecting a consistent non-zero phase lag between neighbouring populations (Fig. 2 A).

The direction and velocity of the travelling waves was quantified using nodal phase velocity vectors, which index the average phase relationship between each local population and its neighbours (Roberts et al., 2019). The transition from global synchrony (Fig. 2 C) to travelling waves is accompanied by a corresponding transition to a common alignment of phase vectors in the direction of the wave propagation (Fig. 2E, G and Suppl. Video 3). In the presence of the same asymmetric external current, but flipped in the opposite direction (*p*_head_*> p*_tail_), the wave fronts for slow theta again traverse up the current gradient, hence propagating P-A from the tail to the head (Suppl. Video 4). This is evident in the reverse pattern of phase progression (Fig. 2 F) and a corresponding flip in the direction of the nodal phase vectors (Fig. 2 G).

Despite the reversal of direction, the Kuramoto order parameter approaches a similar value as the waves traversing in the opposite direction (*R* ≈ 0.92 see Fig. 2 A), as it is not sensitive to wave direction, only relative phase alignment.

We next performed a systematic survey of the strength of the current asymmetry for both slow and fast theta, comparing the ensuing estimates against independent empirical data from the literature (Zhang and Jacobs, 2015). In the slow theta regime, asymmetries in external input of ±10% rapidly disrupt global synchrony, with travelling waves emerging within the first second of simulation. This transition is captured by a decline in the Kuramoto order parameter *R* (Fig. 3 A, transition from yellow to blue), indicating the change in phase vectors from strictly coherent to a common spatial offset across the hippocampal surface, in the directions of the travelling waves. Smaller input asymmetries progressively delay the onset of wavefronts, and minimal gradients (when |(Δ*p*)| *<* 2%) fail to support sustained wave propagation (evident as the yellow strip in the centre of the panel). Wavefront velocities for both positive and negative input gradients (Fig. 3 B) fall within the range of travelling waves (*v* = 3 − 5 ms^—1^) observed with depth electrodes during working memory tasks in humans (Zhang and Jacobs, 2015). Notably, velocities of the simulated waves which show more prominent wavefronts (i.e. when |Δ*p*| *>* 4%) lie towards the lower limit observed empirically. Simulated wavelengths also lie within empirical estimates *l* = 0.4 − 1 m, falling slightly below the empirical range for current mismatches of Δ*p >* 4%. The symmetry of these plots indicates that there is no notable difference in velocity or wavelength for waves propagating from the tail to the head compared to the head to the tail of the hippocampus. As noted above, asymmetries in external input also determine wave directionality: for slow theta greater input to the hippocampal head produces P–to-A propagation, while greater input to the tail induces A–P waves (Fig. 3 C). For slow theta, waves thus propagate against the direction of the input current asymmetry (Suppl. Video 4).

**Figure 3:**
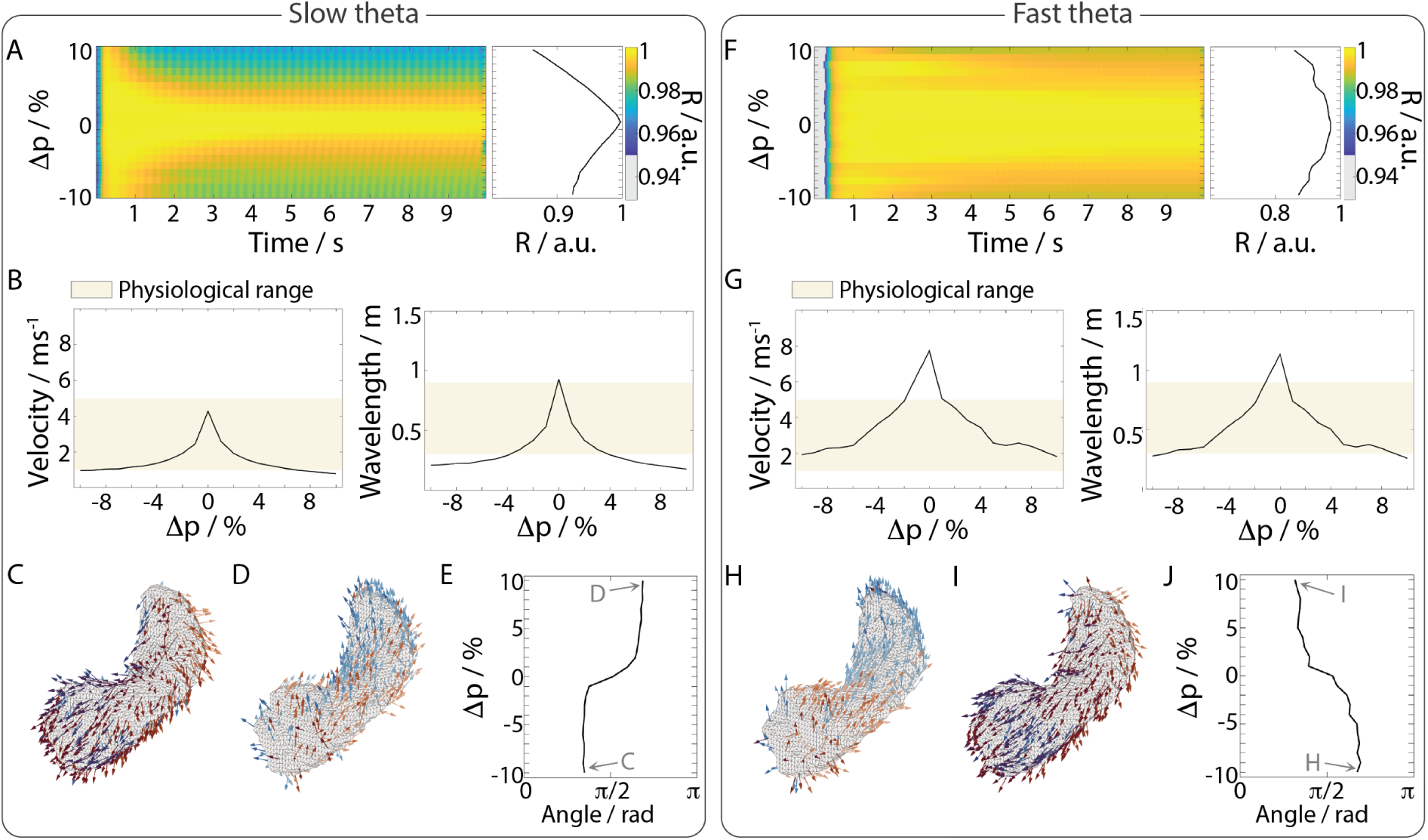
Emergence and behaviour of travelling waves on the hippocampus. A-E. show results for slow theta oscillations, and **F-J** corresponding results for fast theta oscillations. **A, F** Global coherence *R* over time for spatial imbalances in external input *p* between the head and tail of the hippocampus. **B, G** Mean nodal phase velocity and wave-length for different spatial imbalances in external input. The physiological range used for comparison is based on (Zhang and Jacobs, 2015). **C, H** Snapshot of nodal phase vectors for Δ*p* = −10% at *t* = 6s, indicating waves propagating in a posterior-to-anterior (P-A) direction. **D, I** Snapshot of nodal phase vectors for Δ*p* = 10% at *t* = 6s, showing waves propagating in an A-P direction. **E, J** Mean angle of nodal velocity vectors relative to the A-P axis. Angles between 0 and *π/*2 indicate waves aligned with the A-P axis.

A wave-derived partial disruption of synchrony also occurs in the fast theta regime under input asymmetries (Fig. 3 F), although the reduction in *R* is less pronounced. Notably, fast theta waves exhibit faster propagation velocities and longer wavelengths relative to slow theta, hence falling into the upper range of empirical estimates (Fig. 3 G). Moreover, the directionality of wave propagation inverts across frequency regimes: In fast theta, tail-dominant input preferentially generates waves that propagate from tail to head (P-A) and conversely head dominant input generates A-P waves (Fig. 3 H-J). Thus fast theta waves propagate down the input current gradient, in contrast to the slow theta case (Suppl. Video 5).

Modulating the gain of neural systems (changing the slope of the input–output transfer function) is a core computational process (Ermentrout and Kleinfeld, 2001a; Muller et al., 2018; Zhang and Jacobs, 2015; Zhang et al., 2018), mediating the response to, or anticipation of perceptual inputs (Shine et al., 2021). Cortical and ascending neuromodulatory systems regulate hippocampal gain in a topographically organised manner along its A-P axis (Herńandez-Frausto and Vivar, 2024; Maleki Balajoo et al., 2023). The anterior hippocampus receives dense cholinergic, noradrenergic, and dopaminergic inputs from medial prefrontal, orbitofrontal, and limbic cortices that increase neural gain (Amaral and Kurz, 1985; Gasbarri et al., 1994). In contrast, the posterior hippocampus operates in a lower-gain regime via modulating serotonergic projections from retrosplenial and parietal cortices (Michelsen et al., 2007; Vertes, 1991). Together, these neurotransmitter gradients support topographically organised gain through which cortical and subcortical networks can flexibly tune hippocampal responsiveness to inputs (Herńandez-Frausto and Vivar, 2024)

To incorporate this effect into our model, we varied neural gain along the A-P hippocampal axis, similar to the study of external input. This was achieved by linearly varying the maximum slope *r* of the sigmoid-shaped gain function (Equation 9, Methods) which maps the local membrane potential to an output firing rate. The value of *r* at the hippocampal tail was held constant at the default value of *r_tail_* = 0.49 mV^—1^ while the corresponding value at the head was varied between *r*_head_ = 0.25 mV^—1^ (gain lower than tail) to *r*_head_ = 0.74 mV^—1^ (gain higher at head).

As with the variation of external input, an asymmetric neuromodulation of gain elicits travelling waves of activity that traverse the hippocampal long axis (Suppl. Fig. S2). For both slow and fast theta oscillations, the travelling waves traverse from low to high gain, hence from head to tail when *r_head_ < r_tail_* (Suppl. Fig. S2 C,H) and conversely from tail to head when *r*_head_ *> r*_tail_ (Suppl. Fig. S2 D,I). For slow theta, the velocity fall into the physiological range for *r*_head_ *>* 0.45 mV^—1^ but the wavelength is shorter than empirical waves for *r*_head_ *<* 0.5 mV^—1^ and *r*_head_ *>* 0.6 mV^—1^ (Suppl. Fig. S2 B). For fast theta, the velocity and wavelength of these waves fall into the physiological range except for *r*_head_ close to 0.55 mV^—1^ (Suppl. Fig. S2 G).

### 2.3 Impact of hippocampal waves on cortical activity

To study the impact of these hippocampal waves on cortical activity, unidirectional coupling between the two surfaces was specified using a functionally-informed empirical embedding, derived from functional Magnetic Resonance Imaging (fMRI) data acquired while healthy adults undertook a naturalistic memory task (Fig. 4 A,B). In brief, this embedding reflects the task specific modulation of hippocampus to cortex co-variation in Blood Oxygenation Level Dependent (BOLD) signal while participants viewed either familiar or novel news items (Borne et al., 2023). Notably, this functional embedding follows a specific topological pattern, with the principal gradient along the A-P axis of the hippocampus mapping onto a corresponding gradient on the cortex that diverges from several peaks within the default mode regions (precuneus, angular gyrus, anterior midline prefrontal cortex and anterior pole of the temporal lobe) across hetero-modal and association cortex, before converging onto primary sensorimotor regions (including motor, somatosensory and visual cortex). We first simulated interactions between cortex and hippocampus by coupling hippocampus unidirectionally to the cortex. Hippocampal activity was set to exhibit propagating waves by virtue of input asymmetries along the A-P axis. In contrast, the external input to the cortex was kept spatially uniform.

**Figure 4:**
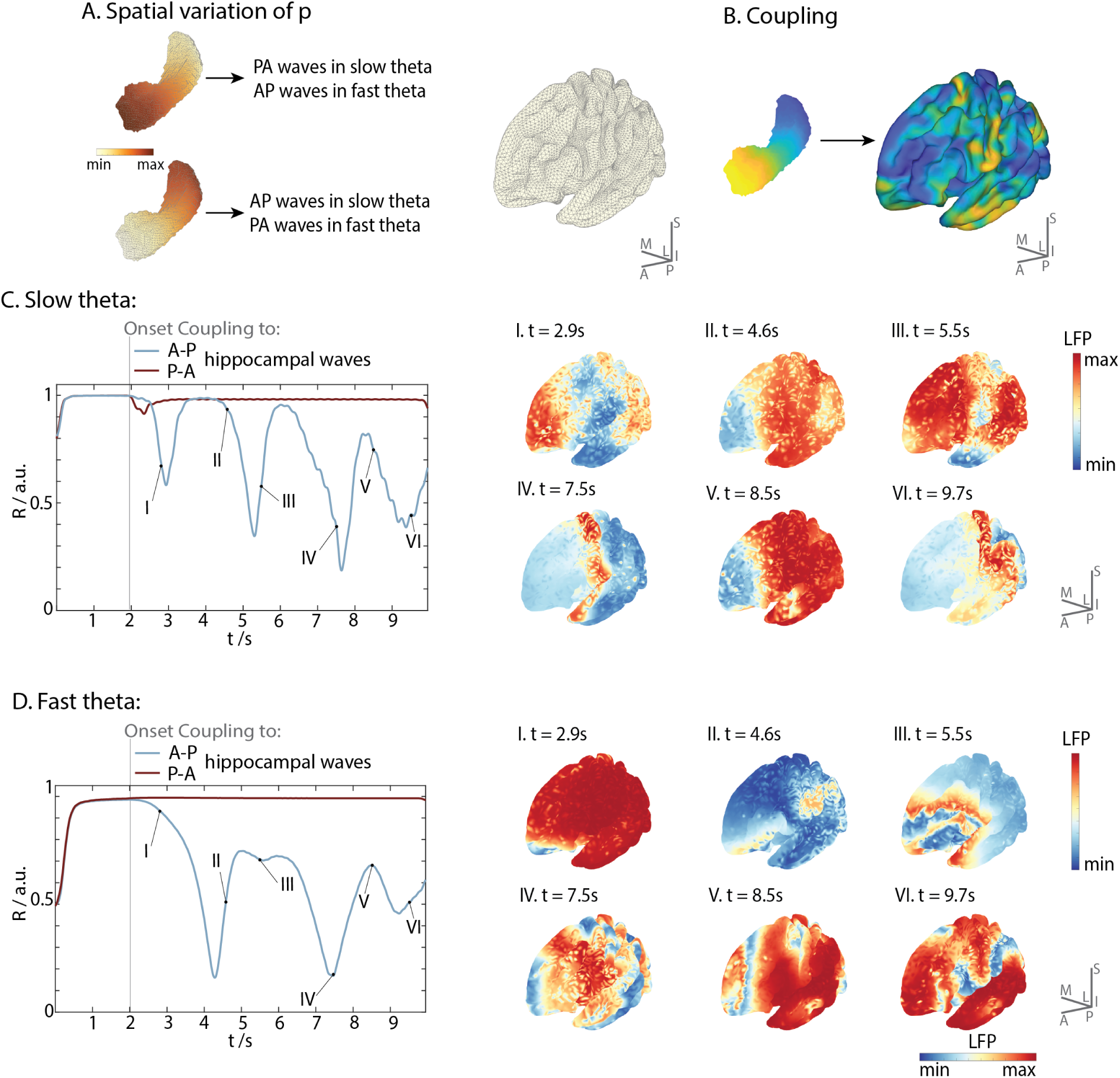
Anterior-to-posterior hippocampal waves reorganise cortical dynamics into travelling waves. (A) Hippocampal input gradient was set to induce hippocampal waves. (B) Hippocampal to cortical coupling was derived from task functional neuroimaging data. Global coherence *R*(*t*) of cortical phase dynamics over time, shown for slow (C) and fast (D) theta regimes. The cortex was coupled from *t* = 2 s onwards to anterior-to-posterior (A–P; blue) and posterior-to-anterior (P–A; red) hippocampal waves. Under coupling from A-P hippocampal waves, in both theta regimes cortical coherence decreases indicative of a transition from global synchrony to spatially structured wave patterns. Cortical reorganisation is reflected in snapshots of the cortical local field potential, revealing transient wavefronts with evolving spatial structure (C, D, right). In contrast, P–A coupling fails to disrupt synchrony with *R*(*t*) remaining near maximal levels.

We allowed the cortex to evolve autonomously for 2 seconds, during which time cortical populations settle onto a synchronous state, as indexed by a high Kuramoto order parameter (Fig. 4 C). For slow theta and hippocampal waves propagating along the A–to-P axis, initiation of the coupling from hippocampus to cortex reorganises cortical dynamics, triggering travelling waves that break the global symmetry and propagate away from one or more spontaneously appearing local sources (Suppl. Video 6 A). Consequently, the Kuramoto order parameter dips well below its maximum value as local phase vectors disperse across the cortical sheet in the outward directions of the waves. Within time scales of 1–2 seconds, the wave fronts from different sources begin to collide along disordered and non stationary fronts, until new sources and sinks appear and the cortex transitions to another global wave state (Fig. 4 C). This metastable switching between sources and wave-to-wave collision is reflected in the waxing and waning of the Kuramoto order parameter (Fig. 4 C).

In contrast to these anterior-to-posterior (A-P) hippocampal waves, hippocampal waves propagating from tail to head (P-A) do not induce travelling wave patterns in the cortex not hence do not disrupt the global cortical synchrony (Fig. 4 C, Suppl. Video 6 B). This situation is mirrored when both hippocampus and cortex exhibit fast theta oscillations - namely A-P waves, but not P-A hippocampal waves, trigger cortical waves that disrupt global synchrony (Fig. 4 D, Suppl. Video 7).

To systematically assess the conditions under which hippocampal activity induces cortical wave propagation, we performed a parameter sweep across a range of asymmetries in the external input Δ*p* and hippocampo-cortical coupling strengths. Across all tested regimes, and for both slow and fast theta, we found that only A-P travelling waves in the hippocampus elicit travelling dynamics in the cortex. In contrast, P-A hippocampal waves failed to re-organise coherent cortical activity into travelling waves, even under strong coupling conditions.

The induced cortical waves are considerably more complex then those on the hippocampus. To characterise the spatial structure of these induced cortical wave patterns, we decomposed the phase flows into their curl and curl-free components (Koller et al., 2024a). The curl-free potential provides a scalar landscape across the cortical surface, with waves propagating from local maxima (sources) toward local minima (sinks). In contrast, the curl component captures the presence and direction of wave rotation during outward and inward propagation. To focus on the appearance and dissolution of sources and sinks, we focused on the curl-free flow potential, using a singular value decomposition (SVD) to quantify the principal wave modes.

In the slow theta regime, the leading spatial mode captures cortical travelling waves propagating posteriorly from a source in prefrontal cortex toward a sink in parietal regions (Fig. 5A), accounting for 78% of the variance. The corresponding temporal mode reveals the onset of structured wave activity following the onset of hippocampal-to-cortical coupling (Fig. 5B). The second spatial mode, explaining 11% of the variance, features sources in frontal and occipital cortices with a sink at the temporal pole. Polarity reversals within this mode, evident as zero crossings in the corresponding amplitude time series, reflect dynamic switching of source–to-sink configurations. The third mode accounts for an additional 3% of the variance and exhibits more spatially fragmented patterns. In the fast theta regime, the leading spatial mode again captures a travelling wave propagating from a frontal source to a posterior sink (Fig. 5C), explaining 64% of the variance. Temporal dynamics capture the reorganisation of cortical dynamics following the onset of hippocampal-cortical coupling (Fig. 5D). The second and third modes explain 24% and 2% of the variance, respectively. As with slow theta, zero crossings of these modes capture metastable switching of sources to sinks and vice versa.

**Figure 5:**
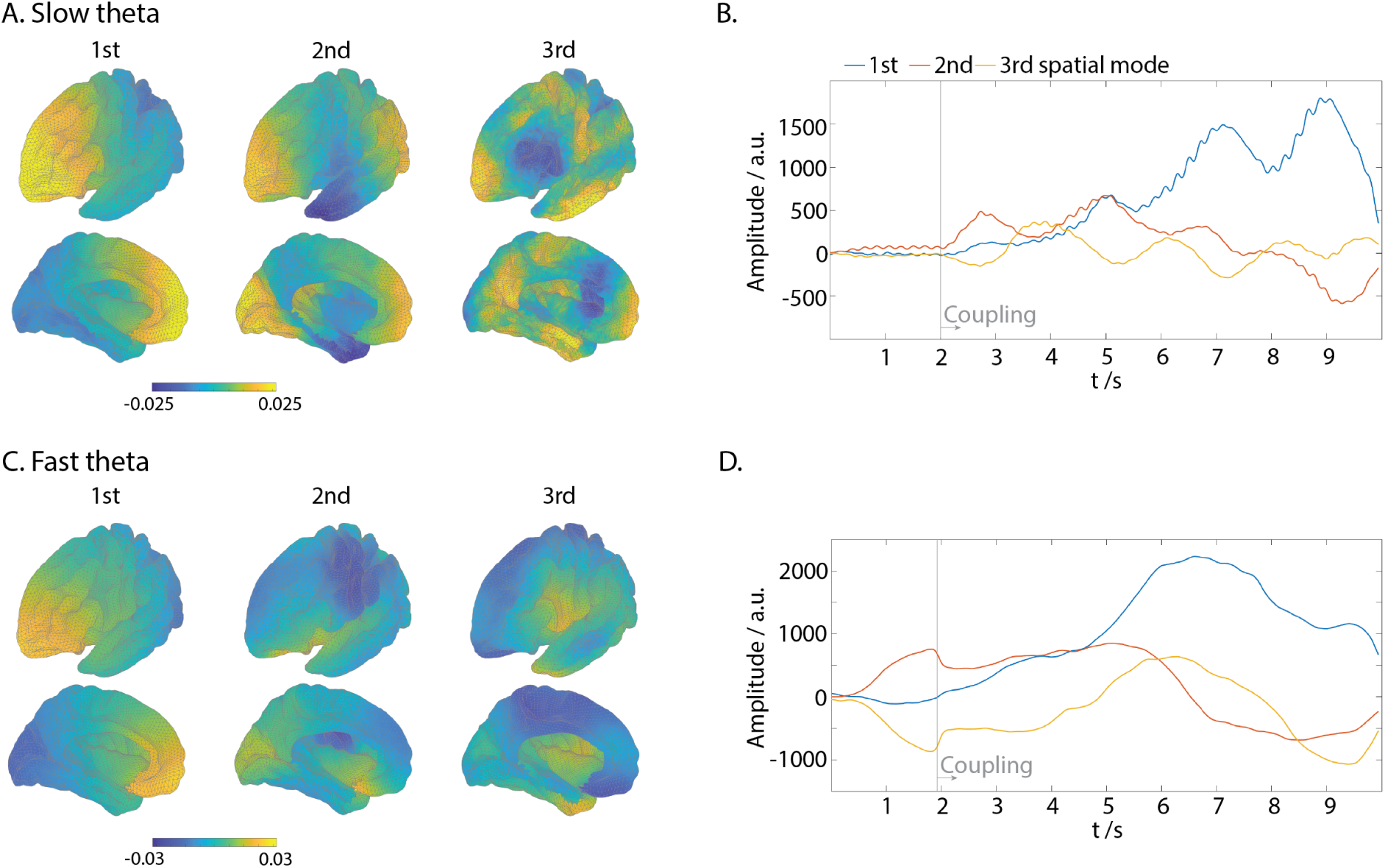
Hippocampal travelling waves induce distinct spatiotemporal patterns of cortical activity. First three spatial modes derived from singular value decomposition (SVD) of the curl-free flow potential on the cortical surface, for **A** slow and **C** fast theta regimes. Waves are directed from regions of higher to lower potential. However, sinks and sources can switch, indicated by changes of sign of the corresponding temporal modes. **B, D** Temporal evolution of the corresponding spatial mode amplitudes reveals amplification of all modes following the onset of hippocampo-cortical coupling at *t* = 2 s. This reflects a transition from baseline synchrony to structured wave dynamics.

### 2.4 Impact of cortical waves on hippocampal activity

We next studied the influence of cortical waves on hippocampal activity. This was achieved by reversing the direction of coupling so that cortical regions projected unidirectionally to hippocampus, via the same empirically derived corticohippocampal coupling matrix (Fig. 4B). The input current and gain within the hippocampus were set to be uniform so that, as above, uncoupled hippocampal activity converges toward global synchrony.

To break the parameter uniformity across the cortex, we introduced a current that scaled according to the principal gradient of cortico-cortical functional connectivity (Margulies et al., 2016). This gradient stretches from one extreme across primary sensory and motor regions, through heteromodal cortex, to peaks in the default-mode network, capturing a functional spectrum from sensory processing and motor action to interoception, mind wandering and introspection (Fig. 6 A). Whereas the mapping between hippocampus and cortex is informed by functional corticohippocampal connectivity, the cortical gradient rests upon a purely cortico-cortical derivation (Margulies et al., 2016). The input current was either scaled linearly from a minimum in default cortex and graded to a maximum across sensorimotor cortex (Fig. 6 A) or reversed and scaled to be minimum across sensorimotor regions grading up to a maximum at the default ”peaks” (Fig. 6 H).

**Figure 6:**
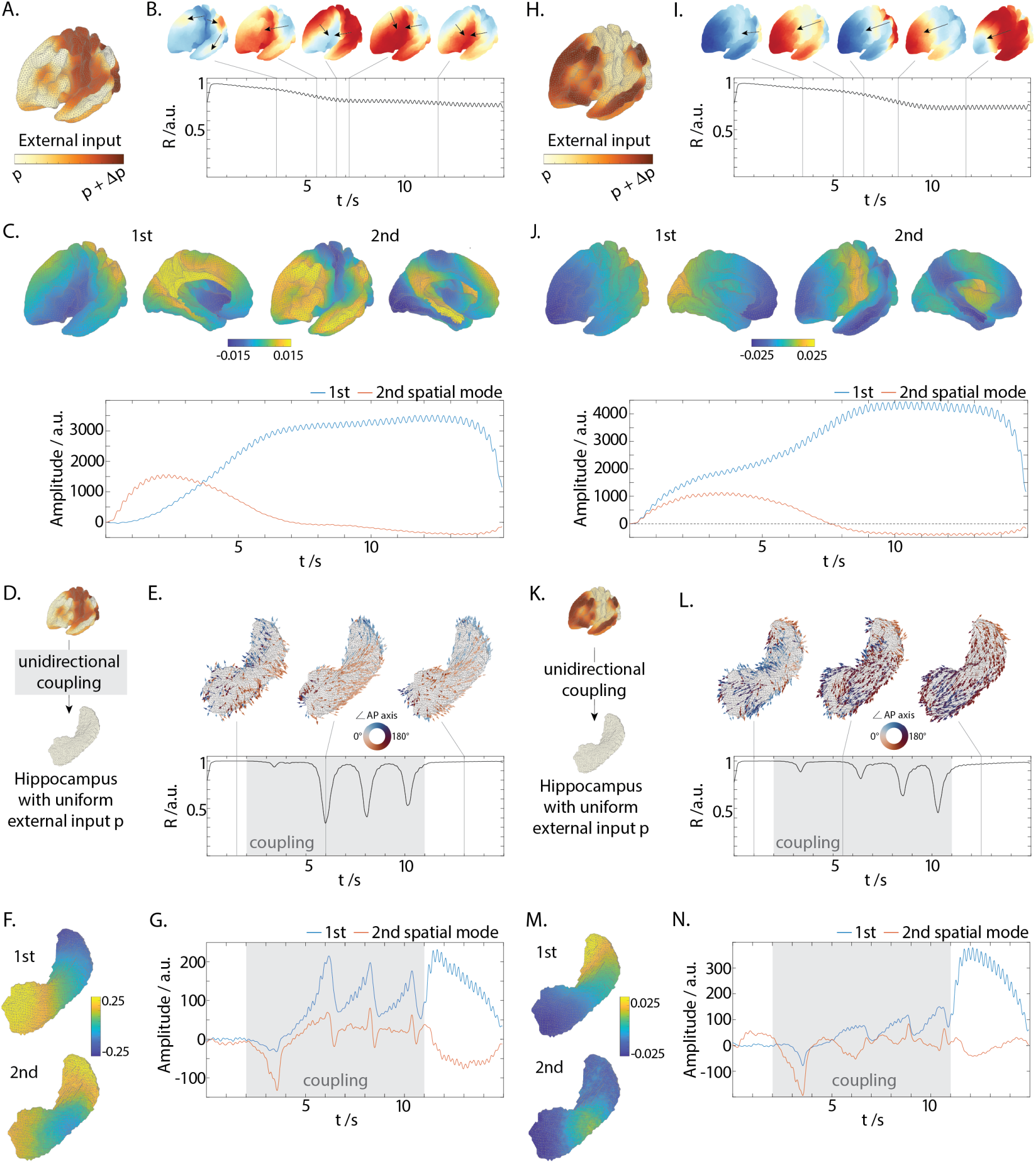
Cortical travelling waves induce travelling waves in the hippocampus. (A) Input current asymmetry minimum in default cortex and maximum across sensorimotor cortex. (B) Propagating and colliding waves disrupt global cortical synchrony and decrease the Kuramoto parameter (C) First and second principle cortical modes from the curl-free flow and their temporal evolution. (D) These cortical waves trigger the intermittent expression of waves from the head to the tail of the hippocampus (E). (F,G) First and second hippocampal modes from the curl-free flow and their temporal evolution. (H) Input current asymmetry minimum in sensorimotor cortex and maximum in default cortex. Corresponding cortical waves (I), their spatial and temporal expression (J) which trigger intermittent waves on the hippocampus (K,L) which propagate from the tail to the head (M,N). Driving hippocampal activity with these slow theta cortical waves triggers hippocampal waves that predominantly traverse from head to tail of the hippocampus (Fig. 6 D,E and Suppl. Video 9). This is reflected in the modal decompositions of the phase flow, which show a dominant head-to-tail bipolar mode whose temporal dynamics capture the waxing and waning evoked waves (Fig. 6 F, G). Notably, a second mode is weakly-co-expressed with the first mode, reflecting a subtle second order influence of cortex on hippocampal waves. Whereas the driving cortical waves are persistent, the hippocampal waves appear intermittently, as captured in sudden dips in the Kuramoto order parameter (Fig. 6 E).

For slow theta, adding a modest input current asymmetry (Δ*p <* 2%) across the cortical gradient elicits the rapid onset of cortical wave fronts. As with slow theta waves on the hippocampus, slow theta waves on the cortex propagate up the input current. Hence, when an input current asymmetry of Δ*p* = 5% is set to be minimum in default cortex and maximum across sensorimotor cortex (Fig. 6 A), cortical waves propagate outwards from the main peaks in default cortex, colliding as they converge towards the central sulcus (Fig. 6 B and Suppl. Video 8). After ≈ 5 seconds, a long-wavelength transition occurs, such that waves originate from a single wave source in the precuneus and propagate across the entire medial and lateral cortex. This is reflected in the time evolution of the principle spatial modes of the flow potential (Fig. 6 C). Expression of the second mode, characterised by multiple peaks in the default mode, peaks at 2 s. The first, whole brain mode then becomes dominate from 4–5 s for the remainder of the run.

Reversing the cortical current asymmetry such that it is minimum across sensorimotor regions and maximum at default cortex yields cortical waves that initially propagate anteriorly from visual cortex and outward from the sensorimotor strip (Fig. 6 H,I). Again at ≈ 5 secs there is a long wavelength transition such that cortical waves propagate from a single source in visual cortex, anteriorly across both medial and lateral walls (Fig. 6 J and Suppl. Video 10). These cortical waves trigger hippocampal waves that traverse from tail to head (Fig. 6 K, L and Suppl. Video 11). Again, these wave patterns are reflecting in the spatial modes estimated on the hippocampal surface which are dominated by a single full length bipolar mode, modulated by a second mode (Fig. 6 M,N).

To study the robustness of these waves, we conducted several parameter surveys: Incremental increases in the corticohippocampal coupling strength led to progressively more frequent discrete wave packets and associated dips in the order parameter (Suppl. Fig. S3). The hippocampal waves are otherwise unchanged, propagating A-P or P-A depending on the direction of the cortical waves (Suppl. Video 12). Cortical and hippocampal waves are robust to variations in the exponent *λ* of the exponential distance rule. Higher values of *λ* (from 0.30 mm^—1^ to 0.60 mm^—1^) correspond to shorter range connectivity and yield waves with shorter wavelengths but their propagation and interactions are otherwise similar to those arising with default settings of *λ*. These patterns are also robust to modest changes in the magnitude of the cortical input asymmetry, with similar cortical and induced hippocampal waves from Δ*p* = 5% − 12%. For greater input asymmetry, Δ*p >* 16%, the cortical waves attain an increasingly strong spiral (curl) component. The sources also display movement away from their initial ”anchors” in default mode peaks, leading to complex and highly non-stationary spatiotemporal mixing (Suppl. Videos only). These patterns in turn elicit more complex hippocampal waves, including sudden switches in the direction of the waves along the long axis of the hippocampus; ”split waves” propagating into or out from a source located in mid hippocampus; and wave fronts that propagate in the transverse direction of the long axis.

## 3 Discussion

Theta travelling waves are a fundamental feature of the hippocampal formation (Lubenov and Siapas, 2009; Patel et al., 2012), but their origins and interactions with cortex are not well understood. Using a biophysically grounded neural mass model, we show that spatial gradients in input or excitability along the hippocampal long axis are sufficient to generate organised travelling waves. When coupled to the cerebral cortex, A-P hippocampal waves consistently reconfigure cortical dynamics, transforming globally synchronous activity into metastable travelling wave patterns. In contrast, P-A waves fail to perturb cortical synchrony, revealing a marked directional asymmetry in cortico-hippocampal communication. Conversely, intrinsically generated cortical waves, arising from their own spatial asymmetries, induce intermittent hippocampal waves, again in a directionally selective manner. Together, these findings identify gradient-constrained wave propagation as a unifying principle governing direction-specific hippocampal–cortical interactions.

These findings position the hippocampus as analogous to a “wave machine” in the brain, capable of generating structured dynamics that propagate across cortical networks, which could transform memory-related hippocampal processes into large scale brain activity. This frames the hippocampus as a low-dimensional generator of structured dynamics, capable of entraining distributed cortical assemblies and embedding episodic content into large-scale neural activity. The complementary directional relationship between relatively simple hippocampal waves and complex, metastable waves on the cortex aligns with theoretical frameworks proposing that memory encoding involves compression of high dimensional cortical activity into low-dimensional hippocampal trajectories, while memory recall reflects the re-expansion of these trajectories back into the cortex (Kerŕen et al., 2025). Wave-based coupling provides a plausible mechanism for this direction-specific transformation.

Theta oscillations in the human medial temporal lobe, including the hippocampus, are crucial for episodic memory encoding and retrieval (Herweg et al., 2020). However, due to its deep location, *in vivo* observation of hippocampal travelling waves is technically challenging. Despite this, longitudinally traversing waves have been observed during working memory encoding (Zhang and Jacobs, 2015) and language processing (Kleen et al., 2021). Our model offers a biophysical explanation for these observations, showing how spatial asymmetries in input and excitability along the hippocampal axis are sufficient to generate longitudinal travelling waves. It also reproduces the empirical finding that wavefronts in the fast theta regime propagate faster and with longer wavelengths than in slow theta (Kleen et al., 2021; Zhang and Jacobs, 2015). Importantly, our findings suggest that wave directionality alone does not define functional state. Instead, it is the interplay between oscillatory regime, input and the spatial profile of gain modulation that sets the cognitive context. For example, fast theta A–P waves may support stimulus-driven encoding and spatial orienting, whereas slow theta A–P waves may reflect top-down reactivation during recall (Goyal et al., 2020; Lega et al., 2012; Miller et al., 2018). The same directionality of hippocampal waves can emerge from different input configurations, consistent with observations that projections from the default mode network onto the hippocampus are observed across distinct cognitive states (Nordin et al., 2025). These spatially organised gradients may underlie the hippocampus’s flexible engagement across cognitive tasks and contribute to the differential vulnerability of memory systems to ageing and disease.

Only anterior-to-posterior (A–P) hippocampal waves successfully induced cortical travelling waves in both fast and slow theta regimes. However, the underlying excitability gradients required to generate these waves differed: in fast theta, A–P waves emerged when anterior (head) regions received greater input, while in slow theta, the same directionality arose from increased posterior input. This shows that wave direction is not simply a by-product of input of gain asymmetry. Only A–P hippocampal waves reorganised cortical dynamics, inducing metastable travelling wave flows across the cortical sheet. This directionality-dependent effect provides a dynamical explanation for how hippocampal state transitions can drive large-scale changes in brain activity. It supports the view that theta wave direction reflects functional state transitions (Kleen et al., 2021), and supports observations that cortical waves typically traverse posterior to anterior during memory encoding and reverse during recall (Mohan et al., 2024).

We employed a task-derived functional embedding of the hippocampus to couple hippocampus and cortex (Borne et al., 2023). This embedding aligns with large-scale cortical gradients which traverse from unimodal to transmodal areas (Vos De Wael et al., 2018), but additionally co-aligns the posterior-to-anterior hippocampal gradient. This topography likely reflects a ”scene-to-self” gradient, where posterior hippocampal regions interface with visual and sensory cortices, and anterior regions map onto the default mode and integrative areas. These patterns mirror macroscale cortical motifs observed in comparative neuroanatomy with anterior hippocampal connectivity targeting heteromodal and paralimbic regions, while posterior fields connect to evolutionarily older sensory areas (Eichert et al., 2024). These spatially organised gradients may underlie the hippocampus’s flexible engagement across cognitive tasks and contribute to the differential vulnerability of memory systems to ageing and disease.

Several limitations of the present work warrant consideration. First, coupling within the hippocampus was approximated using an isotropic, distance-dependent rule, providing a first order description of anatomical constraints (Pang et al., 2023). While this yields a spatially grounded and biologically plausible model of communication, it lacks the specificity afforded by tractography or tracer-based estimates of connectivity. Although recent advances have begun to resolve finer-grained hippocampal subfield organisation (Dalton et al., 2022; Maller et al., 2019), available structural connectomes remain insufficiently resolved to support the vertex-level modelling adopted here. We therefore employed a uniform coupling framework across hippocampus and cortex, enabling direct wave-based interactions while minimising assumptions not directly constrained by current data. Future work could incorporate subfield- and layer-specific connectivity to enhance anatomical realism. Second, gradient-induced wave formation was implemented via spatial variation in external input or gain. Alternative mechanisms, such as spatial gradients in intrinsic frequency or time constants, may also generate structured wave dynamics (Koller et al., 2024b) and are consistent with known physiological heterogeneity along the hippocampal axis (Goyal et al., 2020).

Notably, we examined hippocampus-to-cortex and cortex-to-hippocampus influences in separate unidirectional simulations. The symmetric reversal of unidirectional coupling from cortex to hippocampus and conversely from hippocampus to cortex may be a reasonable approximation for projections that enter and exit via the entorhinal cortex. Incorporating the alternative hippocampal outflow that traverses the fornix would be require an additional, asymmetric matrix (Jones and Breakspear, 2026). In addition, cortex and hippocampus function through bidirectional interactions that are likely critical for processes such as memory consolidation, predictive coding, and perceptual inference. Extending the model to include bidirectional interactions will be important for understanding how ongoing feedback shapes the emergence, stability, and functional role of large-scale travelling waves in both structures.

## 4 Methods

### 4.1 Neural Mass Models

Neuronal activity in a network of neural masses was simulated with a modified version of the original Jansen-Rit model (Jansen and Rit, 1995). The evolution of the average postsynaptic potential (PSP) in each column of neuronal tissue is modulated by three interacting neural populations: pyramidal cells (*y*_0_) with projections to excitatory (*y*_1_) and inhibitory (*y*_2_) interneurons, which both project back to the pyramidal cells. In addition to those excitatory and inhibitory feedback loops, long-range pyramidal-to-pyramidal and short-range inhibitory-to-excitatory interneurons couplings were added to extend the original model and describe macroscale dynamics as shown in Fig. 1 A (Coronel-Oliveros et al., 2021).

The dynamic behaviour of the population average PSP and their temporal derivatives is captured through eight ordinary differential equations, where the first pair of equations describe the influence of pyramidal cells on both local excitatory and inhibitory interneurons, the second pair describes local excitatory input the pyramidal neurons receive, and the third pair describe local inhibitory activity. The combined set of equations for a given node *i* at time *t* are,

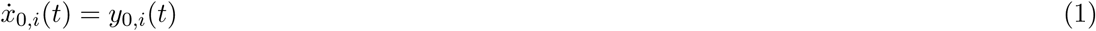

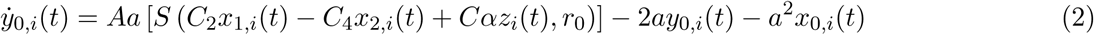

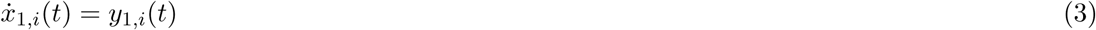

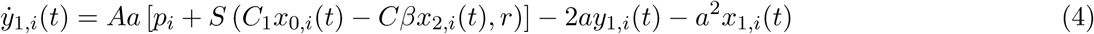

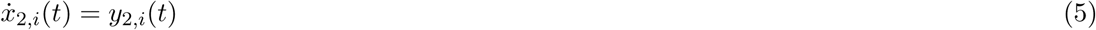

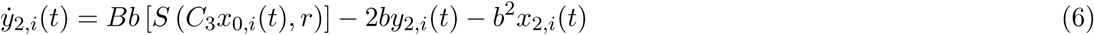

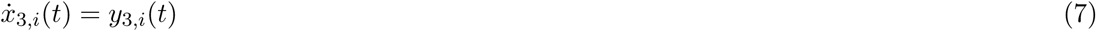

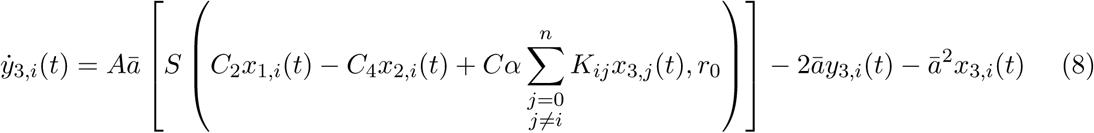

The outputs of pyramidal neurons, excitatory and inhibitory neurons are notated as *x*_0_, *x*_1_ and *x*_2_, and the long-range outputs of pyramidal neurons as *x*_3_. Scaling and time scale paremeters for fast and slow theta are given in (Tab. 1). The connectivity between neural populations is scaled by the constants *C*_1_…_4_. The transduction of neural membrane voltage into a firing rate is given by a sigmoid function of the following form,

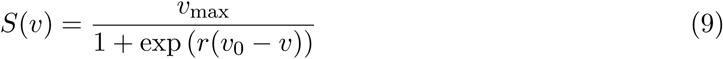

where *v_max_* is the maximum firing rate and *v*_0_ is the average firing threshold of the population. The constant, *r* is the maximum slope of the sigmoid function, centred at the population threshold *v*_0_ which is varied in simulations involving the modulation of neural gain.

### 4.2 Spatial embedding and coupling architecture

#### Surface meshes

The surface mesh of the left hippocampus was generated from the FreeSurfer fsaverage template brain (Fischl et al., 1999), resulting in a triangular mesh with 3,718 vertices. For the cortical representation, the left pial surface of fsaverage5 at 10k resolution was used. To exclude non-functional vertices in the cortical mesh, a medial wall mask was applied. The resulting cortical mesh contained 9,204 vertices.

#### Geodesic distances and within-system coupling

Geodesic distances on both surface meshes were computed using the pygeodesic Python library (version 0.1.9). These distances were used to define the within-system coupling kernel and to derive spatially varying model parameters.

The coupling weight *K_ij_*between Jansen–Rit columns within each system, namely the hippocampus and the cortex, was assumed to follow an EDR,

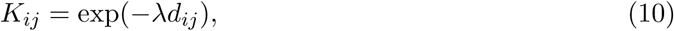

where *d_ij_* is the geodesic distance between nodes *i* and *j*. The decay parameter *λ* varied between 0.15 mm^—1^ and 0.30 mm^—1^ in both systems, consistent with empirical estimates from tracer (Horváat et al., 2016) and diffusion MRI (Roberts et al., 2017) data.

#### Between-system coupling

A sparse one-to-one matrix representing coupling between the hippocampus and the cortex was derived from the projection of the first eigenmap of the hippocampal–cortical functional gradient onto the cortical surface (Borne et al., 2023). This gradient was estimated from fMRI data acquired during naturalistic memory tasks; here, we used the projection corresponding to the continuous memory task in the healthy control group. To construct the coupling matrix, the hippocampal gradient eigenmap and its projection onto grey matter were first transformed from voxel space onto the hippocampal and cortical surfaces using Nilearn (Abraham et al., 2014). Vertices of both meshes were then sorted according to their gradient values, and a sparse one-to-one mapping was obtained by proportionally assigning each hippocampal vertex to a cortical vertex based on its ranked position. This procedure yielded a coupling matrix that preserved correspondences between hippocampal and cortical regions along the shared functional gradient (Fig. 1C).

The resulting matrix defined the topographic pattern of between-system coupling, but not its overall strength. We therefore introduced separate scalar coupling gains for the two directions of interaction: *g*_HC_ for hippocampus-to-cortex coupling and *g*_CH_ for cortex-to-hippocampus coupling. These gains multiplied the activity transmitted across the gradient-informed coupling matrix while preserving its sparse one-to-one structure.

For cortex-to-hippocampus coupling, the additional input to hippocampal node *i* was defined as

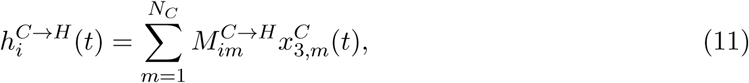

where *M^C^*^→*H*^ is the sparse cortex-to-hippocampus coupling matrix and *x^C^* (*t*) is the long-range pyramidal output of cortical node *m*. This input entered the hippocampal pyramidal-cell equation as

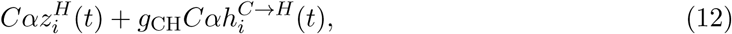

where *z^H^*(*t*) denotes within-hippocampal long-range input. Thus, *g*_CH_ controlled the amplitude of cortical drive to the hippocampus, whereas *C* remained the fixed Jansen–Rit synaptic-contact scaling constant used to define *C*_1_–*C*_4_.

Analogously, for hippocampus-to-cortex coupling, the additional input to cortical node *i* was defined as

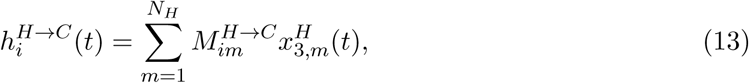

and entered the cortical pyramidal-cell equation as

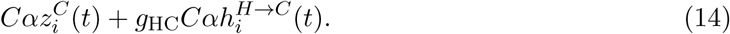

The gradient-informed mapping was therefore identical across simulations, whereas the direction and scalar gain of coupling were varied depending on the experiment. In hippocampus-to-cortex simulations, *g*_HC_ was non-zero and *g*_CH_ = 0. In cortex-to-hippocampus simulations, *g*_CH_ was non-zero and *g*_HC_ = 0. For the parameter sweep shown in Suppl. Fig. S3, *g*_CH_ was systematically varied while hippocampal input *p* and gain *r* were kept spatially uniform.

### 4.3 Parameterisation and spatial tuning

The neural mass models was parameterised using two parameter sets generating distinct theta regimes corresponding to slower (4–7 Hz) and faster (5–9 Hz) oscillatory dynamics. The corresponding model parameters are summarised in Table 1. The average numbers of synaptic contacts between population types were fixed at *C*_1_ = *C* · 1.0, *C*_2_ = *C* · 0.8, *C*_3_ = *C* · 0.25, and *C*_4_ = *C* · 0.25, with *C* = 135.

#### Spatial tuning of hippocampal parameters

To break parameter uniformity across the hippocampus and simulate the influence of topographically organised inputs and neuromodulation, the external stimulus *p* to pyramidal neurons and the sigmoid slope *r* were spatially tuned (Fig. 1A). The values of *p* and *r* at each node were defined as a function of geodesic distance from a representative vertex at the extreme end of the hippocampal tail. The resulting input or gain asymmetry was linearly scaled between the extreme vertices and added to the baseline parameter setting.

#### Spatial tuning of cortical parameters

To break parameter uniformity across the cortex and simulate the influence of topographically organised inputs, the external stimulus *p* to cortical pyramidal neurons was spatially tuned (Fig. 6 A,H). Cortical vertices were rank-ordered according to their scalar value on the principal cortico-cortical gradient described by (Margulies et al., 2016). Input asymmetry was then linearly scaled between the minimum and maximum vertices and added to the baseline parameter setting. This asymmetry was applied either from the default-mode peak vertex (Fig. 6 A) or from the sensorimotor peak vertex (Fig. 6 H), depending on the directionality examined.

### 4.4 Simulations and model outputs

Simulations lasted up to 15 s and were integrated with a time step of 1 ms. To allow the system to stabilise, an equilibration period of 2 s was included during which hippocampal–cortical coupling was inactive. Coupling between hippocampus and cortex was activated at *t* = 2 s and remained active until *t* = 11 s.

Simulated local field potentials (LFPs) were computed as a weighted sum of population activities:

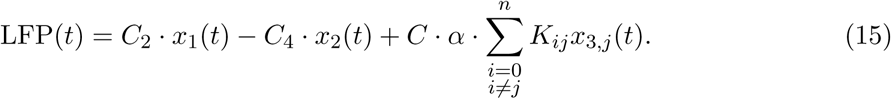

### 4.5 Characterisation of waves

#### Global coherence

Instantaneous phase *φ_j_*(*t*) was extracted at each node *j* using the Hilbert transform. As a quantitative indicator of wave behaviour on the hippocampal or cortical surface, global coherence *R*(*t*) was calculated as

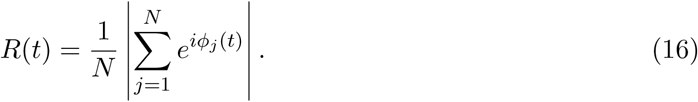

The magnitude of *R* reflects the degree of phase coherence among nodes, ranging from 0 (incoherence) to 1 (perfect synchronisation). When travelling waves are present, *R* typically takes low-to-moderate values.

#### Velocity

The velocity vector field was calculated from the propagation of contours of constant phase, with the direction perpendicular to these contours (Roberts et al., 2019; Rubino et al., 2006). At each node, velocity *→v* was computed from the instantaneous phase *φ*(*x, y, z, t*) as

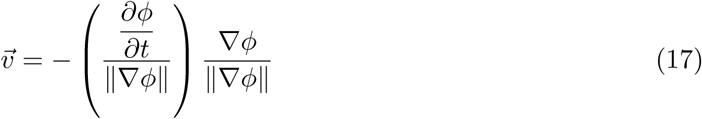

Spatial derivatives were computed using the constrained natural element method (Illoul and Lorong, 2011), which enables calculus operations on non-convex domains. Phase unwrapping was handled using the identity

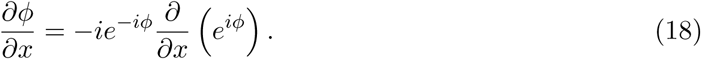

#### Sinks and sources

Wave flow potential was used to characterise wave patterns on the cortical mesh. This scalar field defines a landscape on the cortical surface in which waves can be interpreted as travelling from regions of higher potential (sources) to regions of lower potential (sinks). Flow potential was computed following the workflow described by (Koller et al., 2024b). In brief, the Helmholtz–Hodge decomposition states that a vector field **V**, such as the spatial phase gradient, can be written as

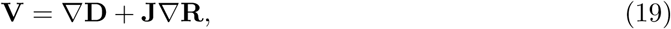

where **D** is the curl-free potential (wave flow potential), **R** is the divergence-free potential, and **J** is a rotation operator (Bhatia et al., 2013). The scalar field **D** was obtained by solving the Poisson equation

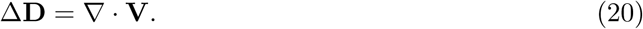

The flow potential was calculated on the cortical mesh for each time point of the simulation. To extract dominant patterns of sources and sinks and assess their contribution over time, a Singular Value Decomposition (SVD) was performed.

## Acknowledgements

The authors acknowledge Dr Nikitas Koussis for assistance with hippocampal mesh extraction.

## Author roles

AB: Methodology, Software, Formal analysis, Writing - Original Draft, RP: Writing - Review & Editing, DTJ: Writing - Review & Editing, JMS: Writing - Review & Editing, MB: Conceptualization, Methodology, Writing - Review & Editing

## References

1. Abraham, A., Pedregosa, F., Eickenberg, M., Gervais, P., Mueller, A., Kossaifi, J., Gramfort, A., Thirion, B., and Varoquaux, G. (2014). Machine learning for neuroimaging with scikit-learn. Frontiers in neuroinformatics, 8:14.

2. Amaral, D. and Kurz, J. (1985). An analysis of the origins of the cholinergic and noncholinergic septal projections to the hippocampal formation of the rat. Journal of Comparative Neurology, 240(1):37–59.

3. Barnett, A. J., Reilly, W., Dimsdale-Zucker, H. R., Mizrak, E., Reagh, Z., and Ranganath, C. (2021). Intrinsic connectivity reveals functionally distinct cortico-hippocampal networks in the human brain. PLOS Biology, 19(6):e3001275.

4. Bhatia, H., Norgard, G., Pascucci, V., and Bremer, P.-T. (2013). The Helmholtz-Hodge Decomposition—A Survey. IEEE Transactions on Visualization and Computer Graphics, 19(8):1386–1404.

5. Borne, L., Tian, Y., Lupton, M. K., Van Der Meer, J. N., Jeganathan, J., Paton, B., Koussis, N., Guo, C. C., Robinson, G. A., Fripp, J., Zalesky, A., and Breakspear, M. (2023). Functional re-organization of hippocampal-cortical gradients during naturalistic memory processes. NeuroImage, 271:119996.

6. Buzsáki, G. (2002). Theta oscillations in the hippocampus. Neuron, 33(3):325–340.

7. Colgin, L. L. (2016). Rhythms of the hippocampal network. Nature Reviews. Neuroscience, 17(4):239–249.

8. Coronel-Oliveros, C., Cofre, R., and Orio, P. (2021). Cholinergic neuromodulation of inhibitory interneurons facilitates functional integration in whole-brain models. PLOS Computational Biology, 17(2):e1008737.

9. Cox, R., Ruber, T., Staresina, B. P., and Fell, J. (2020). Phase-based coordination of hippocampal and neocortical oscillations during human sleep. Communications Biology, 3(1):176.

10. Dalton, M. A., D’Souza, A., Lv, J., and Calamante, F. (2022). New insights into anatomical connectivity along the anterior–posterior axis of the human hippocampus using in vivo quantitative fibre tracking. eLife, 11:e76143.

11. Deco, G., Jirsa, V. K., Robinson, P. A., Breakspear, M., and Friston, K. (2008). The dynamic brain: from spiking neurons to neural masses and cortical fields. PLoS computational biology, 4(8):e1000092.

12. Eichert, N., DeKraker, J., Howard, A. F. D., Huszar, I. N., Zhu, S., Sallet, J., Miller, K. L., Mars, R. B., Jbabdi, S., and Bernhardt, B. C. (2024). Hippocampal connectivity patterns echo macroscale cortical evolution in the primate brain. Nature Communications, 15(1):5963.

13. Ermentrout, G. and Kleinfeld, D. (2001a). Traveling Electrical Waves in Cortex. Neuron, 29(1):33–44.

14. Ermentrout, G. B. and Kleinfeld, D. (2001b). Traveling electrical waves in cortex: insights from phase dynamics and speculation on a computational role. Neuron, 29(1):33–44.

15. Ferris, C., Scheurich, R., Palmer, C., and Sheldon, S. (2025). Hippocampal-cortical networks predict conceptual versus perceptually guided narrative memory. Journal of Neuroscience.

16. Fischl, B., Sereno, M. I., Tootell, R. B., and Dale, A. M. (1999). High-resolution inter-subject averaging and a coordinate system for the cortical surface. Human Brain Mapping, 8(4):272–284.

17. Gasbarri, A., Packard, M. G., Campana, E., and Pacitti, C. (1994). Anterograde and retrograde tracing of projections from the ventral tegmental area to the hippocampal formation in the rat. Brain research bulletin, 33(4):445–452.

18. Genon, S., Bernhardt, B. C., La Joie, R., Amunts, K., and Eickhoff, S. B. (2021). The many dimensions of human hippocampal organization and (dys)function. Trends in Neurosciences, 44(12):977–989.

19. Goyal, A., Miller, J., Qasim, S. E., Watrous, A. J., Zhang, H., Stein, J. M., Inman, C. S., Gross, R. E., Willie, J. T., Lega, B., Lin, J.-J., Sharan, A., Wu, C., Sperling, M. R., Sheth, S. A., McKhann, G. M., Smith, E. H., Schevon, C., and Jacobs, J. (2020). Functionally distinct high and low theta oscillations in the human hippocampus. Nature Communications, 11(1):2469.

20. Hancock, F., Rosas, F. E., Luppi, A. I., Zhang, M., Mediano, P. A., Cabral, J., Deco, G., Kringelbach, M. L., Breakspear, M., Kelso, J. S., et al. (2025). Metastability demystified—the foundational past, the pragmatic present and the promising future. Nature Reviews Neuroscience, 26(2):82–100.

21. Heitmann, S., Boonstra, T., and Breakspear, M. (2013). A dendritic mechanism for decoding traveling waves: principles and applications to motor cortex. PLoS computational biology, 9(10):e1003260.

22. Heitmann, S., Gong, P., and Breakspear, M. (2012). A computational role for bistability and traveling waves in motor cortex. Frontiers in computational neuroscience, 6:67.

23. Hernández-Frausto, M. and Vivar, C. (2024). Entorhinal cortex–hippocampal circuit connectivity in health and disease. Frontiers in human neuroscience, 18:1448791.

24. Herweg, N. A., Solomon, E. A., and Kahana, M. J. (2020). Theta Oscillations in Human Memory. Trends in Cognitive Sciences, 24(3):208–227.

25. Horvát, S., Gămănut, R., Ercsey-Ravasz, M., Magrou, L., Gămănut, B., Van Essen, D. C., Burkhalter, A., Knoblauch, K., Toroczkai, Z., and Kennedy, H. (2016). Spatial Embedding and Wiring Cost Constrain the Functional Layout of the Cortical Network of Rodents and Primates. PLOS Biology, 14(7):e1002512.

26. Illoul, L. and Lorong, P. (2011). On some aspects of the CNEM implementation in 3D in order to simulate high speed machining or shearing. Computers & Structures, 89(11-12):940–958.

27. Jansen, B. H. and Rit, V. G. (1995). Electroencephalogram and visual evoked potential generation in a mathematical model of coupled cortical columns. Biological Cybernetics, 73(4):357–366.

28. Jones, D. and Breakspear, M. (2026). The hippocampal latent diffusion engine: A computational framework for memory, perception, and cognitive dysfunction. PsyArXiv, tnygx_v_1.

29. Kerren, C., Reznik, D., Doeller, C. F., and Griffiths, B. J. (2025). Exploring the role of dimensionality transformation in episodic memory. Trends in Cognitive Sciences, page S136466132500021X.

30. Kleen, J. K., Chung, J. E., Sellers, K. K., Zhou, J., Triplett, M., Lee, K., Tooker, A., Haque, R., and Chang, E. F. (2021). Bidirectional propagation of low frequency oscillations over the human hippocampal surface. Nature Communications, 12(1):2764.

31. Koller, D. P., Schirner, M., and Ritter, P. (2024a). Human connectome topology directs cortical traveling waves and shapes frequency gradients. Nature Communications, 15(1):3570.

32. Koller, D. P., Schirner, M., and Ritter, P. (2024b). Human connectome topology directs cortical traveling waves and shapes frequency gradients. Nature Communications, 15(1):3570.

33. Lee, S., Park, J., Alekseichuk, I., Berger, T. A., Manea, A. M. G., Tran, H., Delgado Salazar, G., Konig, S. D., Herman, A. B., Darrow, D. P., Zimmermann, J., and Opitz, A. (2026). Traveling-wave transcranial alternating current stimulation (twtACS) causally links neural timing to cognitive function. Proceedings of the National Academy of Sciences, 123(19):e2527296123.

34. Lega, B. C., Jacobs, J., and Kahana, M. (2012). Human hippocampal theta oscillations and the formation of episodic memories. Hippocampus, 22(4):748–761.

35. Lubenov, E. V. and Siapas, A. G. (2009). Hippocampal theta oscillations are travelling waves. Nature, 459(7246):534–539.

36. Maleki Balajoo, S., Eickhoff, S. B., Masouleh, S. K., Plachti, A., Waite, L., Saberi, A., Bahri, M. A., Bastin, C., Salmon, E., Hoffstaedter, F., et al. (2023). Hippocampal metabolic subregions and networks: Behavioral, molecular, and pathological aging profiles. Alzheimer’s & Dementia, 19(11):4787–4804.

37. Maller, J. J., Welton, T., Middione, M., Callaghan, F. M., Rosenfeld, J. V., and Grieve, S. M. (2019). Revealing the Hippocampal Connectome through Super-Resolution 1150-Direction Diffusion MRI. Scientific Reports, 9(1):2418.

38. Margulies, D. S., Ghosh, S. S., Goulas, A., Falkiewicz, M., Huntenburg, J. M., Langs, G., Bezgin, G., Eickhoff, S. B., Castellanos, F. X., Petrides, M., et al. (2016). Situating the default-mode network along a principal gradient of macroscale cortical organization. Proceedings of the National Academy of Sciences, 113(44):12574–12579.

39. Massimini, M., Huber, R., Ferrarelli, F., Hill, S., and Tononi, G. (2004). The sleep slow oscillation as a traveling wave. Journal of Neuroscience, 24(31):6862–6870.

40. Michelsen, K. A., Schmitz, C., and Steinbusch, H. W. (2007). The dorsal raphe nucleus—from silver stainings to a role in depression. Brain research reviews, 55(2):329–342.

41. Miller, J., Watrous, A. J., Tsitsiklis, M., Lee, S. A., Sheth, S. A., Schevon, C. A., Smith, E. H., Sperling, M. R., Sharan, A., Asadi-Pooya, A. A., Worrell, G. A., Meisenhelter, S., Inman, C. S., Davis, K. A., Lega, B., Wanda, P. A., Das, S. R., Stein, J. M., Gorniak, R., and Jacobs, J. (2018). Lateralized hippocampal oscillations underlie distinct aspects of human spatial memory and navigation. Nature Communications, 9(1):2423.

42. Mohan, U. R., Zhang, H., Ermentrout, B., and Jacobs, J. (2024). The direction of theta and alpha travelling waves modulates human memory processing. Nature Human Behaviour.

43. Muller, L., Chavane, F., Reynolds, J., and Sejnowski, T. J. (2018). Cortical travelling waves: mechanisms and computational principles. Nature Reviews Neuroscience, 19(5):255–268.

44. Muller, L. and Destexhe, A. (2012). Propagating waves in thalamus, cortex and the thalamocortical system: experiments and models. Journal of Physiology-Paris, 106(5-6):222–238.

45. Muller, L., Piantoni, G., Koller, D., Cash, S. S., Halgren, E., and Sejnowski, T. J. (2016). Rotating waves during human sleep spindles organize global patterns of activity that repeat precisely through the night. eLife, 5:e17267.

46. Naoumenko, D. and Gong, P. (2019). Complex dynamics of propagating waves in a two-dimensional neural field. Frontiers in Computational Neuroscience, 13:50.

47. Nordin, K., Pedersen, R., Falahati, F., Johansson, J., Grill, F., Andersson, M., Korkki, S. M., Backman, L., Zalesky, A., Rieckmann, A., Nyberg, L., and Salami, A. (2025). Two long-axis dimensions of hippocampal-cortical integration support memory function across the adult lifespan. eLife, 13:RP97658.

48. Olsen, R. K., Moses, S. N., Riggs, L., and Ryan, J. D. (2012). The hippocampus supports multiple cognitive processes through relational binding and comparison. Frontiers in human neuroscience, 6:146.

49. Pang, J. C., Aquino, K. M., Oldehinkel, M., Robinson, P. A., Fulcher, B. D., Breakspear, M., and Fornito, A. (2023). Geometric constraints on human brain function. Nature, 618(7965):566–574.

50. Paquola, C., Benkarim, O., DeKraker, J., Lariviere, S., Frassle, S., Royer, J., Tavakol, S., Valk, S., Bernasconi, A., Bernasconi, N., Khan, A., Evans, A. C., Razi, A., Smallwood, J., and Bernhardt, B. C. (2020). Convergence of cortical types and functional motifs in the human mesiotemporal lobe. eLife, 9:e60673.

51. Paquola, C., Vos De Wael, R., Wagstyl, K., Bethlehem, R. A., Hong, S.-J., Seidlitz, J., Bullmore, E. T., Evans, A. C., Misic, B., Margulies, D. S., et al. (2019). Microstructural and functional gradients are increasingly dissociated in transmodal cortices. PLoS biology, 17(5):e3000284.

52. Patel, J., Fujisawa, S., Berenyi, A., Royer, S., and Buzsaki, G. (2012). Traveling theta waves along the entire septotemporal axis of the hippocampus. Neuron, 75(3):410–417.

53. Plachti, A., Eickhoff, S. B., Hoffstaedter, F., Patil, K. R., Laird, A. R., Fox, P. T., Amunts, K., and Genon, S. (2019). Multimodal parcellations and extensive behavioral profiling tackling the hippocampus gradient. Cerebral cortex, 29(11):4595–4612.

54. Poppenk, J., Evensmoen, H. R., Moscovitch, M., and Nadel, L. (2013). Long-axis specialization of the human hippocampus. Trends in cognitive sciences, 17(5):230–240.

55. Przézdzik, I., Faber, M., Fernández, G., Beckmann, C. F., and Haak, K. V. (2019). The functional organisation of the hippocampus along its long axis is gradual and predicts recollection. Cortex, 119:324–335.

56. Raut, R. V., Snyder, A. Z., Mitra, A., Yellin, D., Fujii, N., Malach, R., and Raichle, M. E. (2021). Global waves synchronize the brain’s functional systems with fluctuating arousal. Science Advances, 7(30):eabf2709.

57. Roberts, J. A., Gollo, L. L., Abeysuriya, R. G., Roberts, G., Mitchell, P. B., Woolrich, M. W., and Breakspear, M. (2019). Metastable brain waves. Nature Communications, 10(1):1056.

58. Roberts, J. A., Perry, A., Roberts, G., Mitchell, P. B., and Breakspear, M. (2017). Consistency-based thresholding of the human connectome. NeuroImage, 145:118–129.

59. Robinson, P. A., Rennie, C. J., and Wright, J. J. (1997). Propagation and stability of waves of electrical activity in the cerebral cortex. Physical Review E, 56(1):826.

60. Rubino, D., Robbins, K. A., and Hatsopoulos, N. G. (2006). Propagating waves mediate information transfer in the motor cortex. Nature Neuroscience, 9(12):1549–1557.

61. Shine, J. M., Muller, E. J., Munn, B., Cabral, J., Moran, R. J., and Breakspear, M. (2021). Computational models link cellular mechanisms of neuromodulation to large-scale neural dynamics. Nature neuroscience, 24(6):765–776.

62. Vertes, R. P. (1991). A pha-l analysis of ascending projections of the dorsal raphe nucleus in the rat. Journal of Comparative Neurology, 313(4):643–668.

63. Vogel, J. W., La Joie, R., Grothe, M. J., Diaz-Papkovich, A., Doyle, A., Vachon-Presseau, E., Lepage, C., Vos de Wael, R., Thomas, R. A., Iturria-Medina, Y., et al. (2020). A molecular gradient along the longitudinal axis of the human hippocampus informs largescale behavioral systems. Nature communications, 11(1):960.

64. Vos De Wael, R., Lariviere, S., Caldairou, B., Hong, S.-J., Margulies, D. S., Jefferies, E., Bernasconi, A., Smallwood, J., Bernasconi, N., and Bernhardt, B. C. (2018). Anatomical and microstructural determinants of hippocampal subfield functional connectome embedding. Proceedings of the National Academy of Sciences, 115(40):10154–10159.

65. Whittington, J. C., McCaffary, D., Bakermans, J. J., and Behrens, T. E. (2022). How to build a cognitive map. Nature neuroscience, 25(10):1257–1272.

66. Xu, Y., Long, X., Feng, J., and Gong, P. (2023). Interacting spiral wave patterns underlie complex brain dynamics and are related to cognitive processing. Nature human behaviour, 7(7):1196–1215.

67. Zhang, H. and Jacobs, J. (2015). Traveling Theta Waves in the Human Hippocampus. Journal of Neuroscience, 35(36):12477–12487.

68. Zhang, H., Watrous, A. J., Patel, A., and Jacobs, J. (2018). Theta and Alpha Oscillations Are Traveling Waves in the Human Neocortex. Neuron, 98(6):1269–1281.e4.

69. Zheng, A., Montez, D. F., Marek, S., Gilmore, A. W., Newbold, D. J., Laumann, T. O., Kay, B. P., Seider, N. A., Van, A. N., Hampton, J. M., Alexopoulos, D., Schlaggar, B. L., Sylvester, C. M., Greene, D. J., Shimony, J. S., Nelson, S. M., Wig, G. S., Gratton, C., McDermott, K. B., Raichle, M. E., Gordon, E. M., and Dosenbach, N. U. F. (2021a). Parallel hippocampal-parietal circuits for self- and goal-oriented processing. Proceedings of the National Academy of Sciences, 118(34):e2101743118.

70. Zheng, A., Montez, D. F., Marek, S., Gilmore, A. W., Newbold, D. J., Laumann, T. O., Kay, B. P., Seider, N. A., Van, A. N., Hampton, J. M., et al. (2021b). Parallel hippocampal-parietal circuits for self-and goal-oriented processing. Proceedings of the National Academy of Sciences, 118(34):e2101743118.

