## Supplemental Figures for "Gradients in excitability generate hippocampal waves and shape their interactions with cortex"

### Supplementary Material

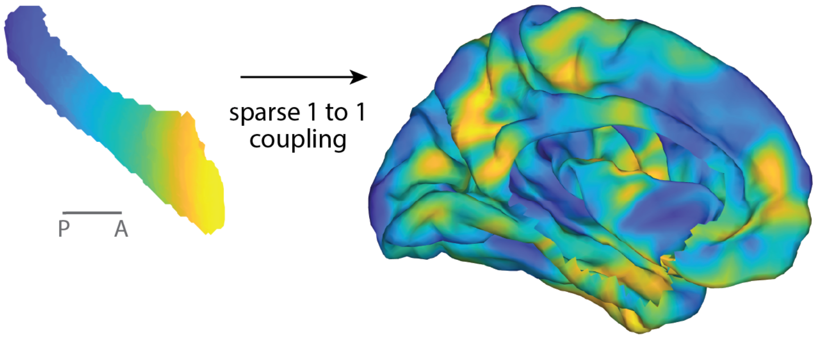


**Figure S1: Lateral view of hippocampus-to-cortex coupling.** Lateral cortical view of the sparse hippocampus-to-cortex coupling scheme used in the model. Each hippocampal vertex was connected to exactly one cortical vertex, producing a one-to-one mapping informed by human functional connectivity data. Matching colours indicate corresponding hippocampal and cortical vertices. This view complements the main figure by showing the spatial distribution of cortical coupling targets on the lateral surface.

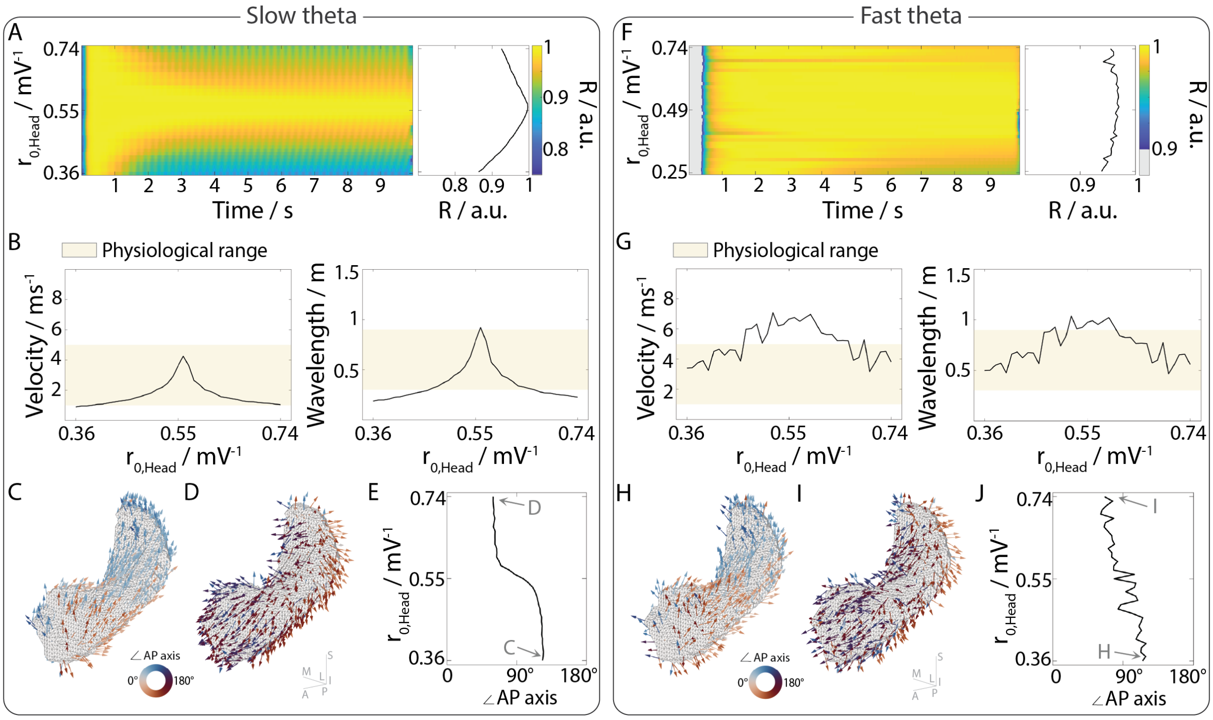


**Figure S2: Emergence and behaviour of travelling waves on the hippocampus with spatially varying gain modulation.** **A-E** show results for slow theta oscillations, and **F-J** corresponding results for fast theta oscillations. **A, F** Global coherence $R$ over time for spatial imbalances in pyramidal gain $r_0$ between the head and tail of the hippocampus. **B, G** Mean nodal phase velocity and wave length for different spatial imbalances in neural gain. $r_{\text{Tail}}$ was held constant at $0.55\,\text{mV}^{-1}$ while $r_{\text{Head}}$ was between $0.36...0.74\,\text{mV}^{-1}$). The physiological range used for comparison is based on (Zhang and Jacobs, 2015). **C, H** Snapshot of nodal phase vectors for $r_{\text{Head}} = 0.36\,\text{mV}^{-1}$) at $t = 6\,\text{s}$, indicating waves propagating in a posterior-to-anterior (P-A) direction. **D, I** Snapshot of nodal phase vectors for $r_{\text{Head}} = 0.74\,\text{mV}^{-1}$) at $t = 6\,\text{s}$, showing waves propagating in an A-P direction. **E, J** Mean angle of nodal velocity vectors relative to the A-P axis. Thereby, 0° indicates perfect alignment with in P to A direction.

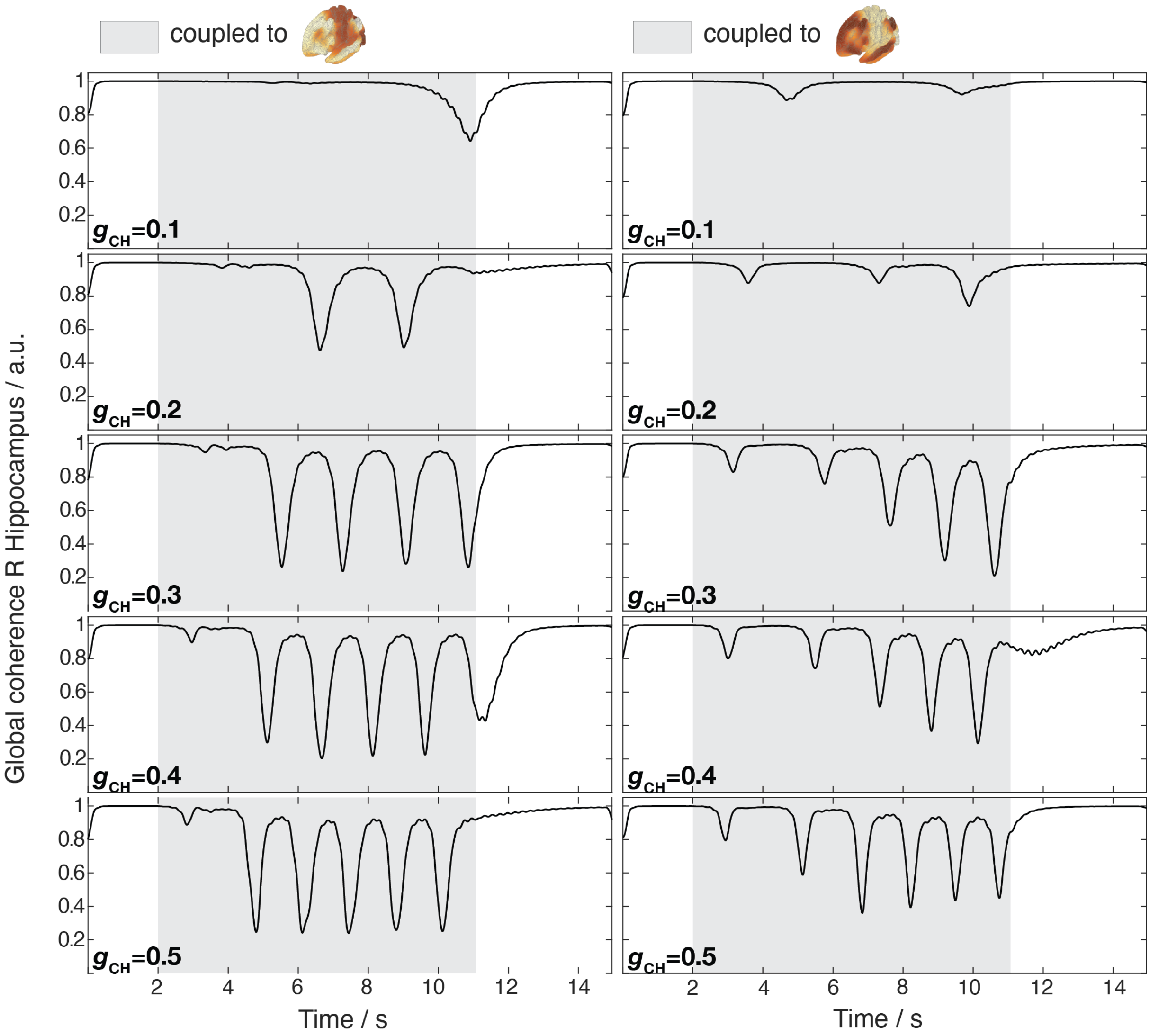


**Figure S3: Cortex-driven hippocampal wave induction depends on cortico-hippocampal coupling strength.** The hippocampal order parameter $R$ is shown over time for two cortical input configurations (columns) and increasing cortico-hippocampal coupling strength $g_{\text{CH}}$ (rows, weakest at top and strongest at bottom). The hippocampal input was spatially uniform in all simulations, so transient deviations in hippocampal dynamics arise from cortical coupling rather than direct hippocampal stimulation. At weak coupling, hippocampal coherence remains close to $R = 1$, with only rare transient reductions. As coupling strength increases, cortical activity induces increasingly frequent drops in $R$, consistent with more frequent hippocampal wave events or transient loss of global phase coherence. The two cortical input configurations differ in the timing and strength of these induced hippocampal events, indicating that both coupling strength and cortical input structure shape hippocampal wave dynamics.
